# rRNA Operon Improves Species-Level Classification of Bacteria and Microbial Community Analysis Compared to 16S rRNA

**DOI:** 10.1101/2024.04.01.587560

**Authors:** Sohyoung Won, Seoae Cho, Heebal Kim

## Abstract

Precise identification of species is fundamental in microbial genomics, crucial for understanding the microbial communities. While the 16S rRNA gene, particularly its V3-V4 regions, has been extensively employed for microbial identification, however has limitations in achieving species-level resolution. Advancements in long-read sequencing technologies have highlighted the rRNA operon as a more accurate marker for microbial classification and analysis than the 16S rRNA gene. This study aims to compare the accuracy of species classification and microbial community analysis using the rRNA operon versus the 16S rRNA gene. We evaluated the species classification accuracy of the rRNA operon,16S rRNA gene, and 16S rRNA V3-V4 region using a BLAST based method and a *k*-mer matching based method with public data available from NCBI. We further preformed simulations to model microbial community analysis. We accessed the performance using each marker in community composition estimation and differential abundance analysis. Our findings demonstrate that the rRNA operon offers an advantage over the 16S rRNA gene and its V3-V4 region for species-level classification within genus. When applied to microbial community analysis, the rRNA operon enables a more accurate determination of composition. Using the rRNA operon yielded more reliable results in differential abundance analysis as well.

**IMPORTANCE:** We quantitatively demonstrated that the rRNA operon outperformed the 16S rRNA and its V3-V4 regions in accuracy, for both individual species identification and species-level microbial community analysis. Our findings can provide guidelines for selecting appropriate markers in the field of microbial research.

## INTRODUCTION

Accurate taxonomic classification is crucial for reliable outcomes in microbial genomics research. As analysis increasingly shifts towards species-level identification beyond the genus level, enhancing the resolution of microbial identification becomes critical for discerning specific species (1). This plays a significant role in discovering novel microbial species and fostering a comprehensive understanding of microbial communities (1).

Next-generation sequencing (NGS) technologies have revolutionized microbial genomics, enabling rapid and cost-effective sequencing of whole genomes and amplicons (2). Second-generation sequencing platforms, like Illumina’s HiSeq and MiSeq, generate millions of short reads (100-300 bp) (3), while third-generation technologies, like PacBio’s SMRT and Oxford Nanopore’s MinION, produce significantly longer reads (up to 100 kb or more) (4, 5).

16S rRNA gene sequencing is a widely used method for microbial identification and community profiling (6, 7). It targets the highly conserved 16S rRNA gene, containing variable regions among species. Some second-generation sequencing approaches using only specific variable regions (e.g., V3 and V4) as markers can be cost-effective but have limitations in taxonomic resolution (8). Even utilizing the entire 16S rRNA gene, accurate species-level classification remains challenging, potentially underestimating diversity and hindering accurate characterization of microbial communities (9, 10).

The emergence of third-generation sequencing technologies has enabled the analysis of larger genomic regions, paving the way for whole rRNA operon sequencing as a prominent approach (11). Encompassing the 16S, 23S, and 5S rRNA genes, along with the Internal Transcribed Spacer (ITS) regions, the rRNA operon provides a comprehensive framework for microbial identification and phylogenetic studies (12). Compared to 16S rRNA sequencing, rRNA operon sequencing offers richer information content, promising higher-resolution taxonomic classification, reaching the species level and more accurate microbiome community analysis (13). However, further quantitative research is required to fully validate these expectations.

This study utilizes public data to compare the accuracy of species classification within the same genus using the entire rRNA operon sequence, the 16S rRNA sequence, and the V3 and V4 regions of the 16S rRNA. Additionally, we create simulated microbiome community data to compare how accurately each region determines the proportion of each species. The aim is to provide guidelines for selecting marker regions for bacterial species classification and species-level microbiome studies.

## RESULTS

### Species classification accuracy within genus

Both BLAST and k-mer matching methods demonstrated significantly higher accuracy when utilizing the entire rRNA operon compared to the 16S rRNA alone (Fig. 1). The average accuracy for BLAST-based classification using the rRNA operon reached 0.999, with a standard deviation of 0.005. This accuracy dropped to 0.936 with a standard deviation of 0.108 when using the 16S rRNA, and further decreased to 0.689 with a standard deviation of 0.300 with the 16S rRNA V3-V4 regions. This trend reflects that analyzing broader genomic regions leads to improved accuracy and reduced variability.

**FIG 1.**
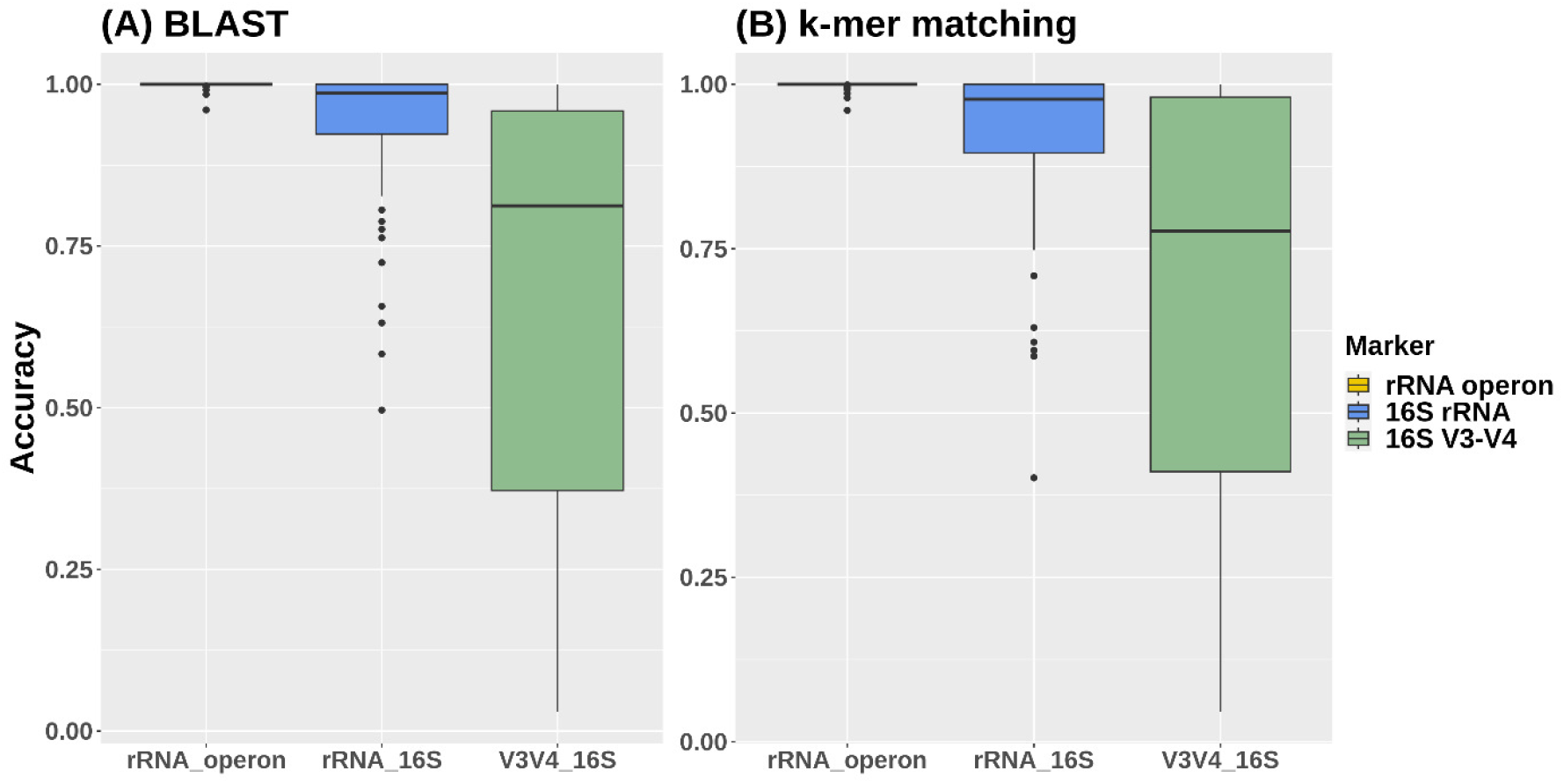
A boxplot of species classification accuracy across genera using the rRNA operon versus the 16S rRNA gene and its V3-V4 regions: (A) demonstrates the results from the BLAST method, showing a median accuracy of 0.999 (SD: 0.005) for the rRNA operon, 0.936 (SD: 0.108) for the 16S rRNA, and 0.689 (SD: 0.300) for the V3-V4 regions; (B) presents outcomes from the k-mer matching method, with a median accuracy of 0.999 (SD: 0.006) for the rRNA operon, 0.918 (SD: 0.123) for the 16S rRNA, and 0.693 (SD: 0.297) for the V3-V4 regions.

k-mer matching yielded comparable results. The average accuracy using the rRNA operon was 0.999, exceeding the 0.918 observed for the 16S rRNA and 0.693 for the V3-V4 regions. The rRNA operon also displayed the lowest standard deviation (0.006), compared to 0.123 for the 16S rRNA and 0.297 for the V3-V4 regions.

Across both methods, the *Haemophilus* genus exhibited the lowest accuracy with the rRNA operon, which was 0.960. For the 16S rRNA, the lowest accuracy was observed in the *Serratia* genus, with BLAST and k-mer matching methods reporting 0.402 and 0.496, respectively. Notably, employing the rRNA operon compared to the 16S rRNA consistently achieved higher accuracy for all genera when using the k-mer matching method. The BLAST method presented a single minor exception in *Chlamydia*, where 16S rRNA yielded marginally higher accuracy with a difference of only 0.0002. On average, the rRNA operon achieved a classification accuracy 0.084 higher with BLAST and 0.109 higher with k-mer matching compared to the 16S rRNA. The largest observed difference in a single genus reached a substantial gap of 0.503 (BLAST) and 0.598 (k-mer matching). Using the rRNA operon, the BLAST method achieved perfect species classification accuracy (1.0) in 89.6% (43) of genera, and the k-mer match method did so in 83.3% (40) of genera. In contrast, with the 16S rRNA, the BLAST method had less than 0.9 accuracy in 31.3% (15) of genera, and the k-mer matching method in 37.5% (18) of genera. This indicates that using the rRNA operon enables more precise species classification than the 16S rRNA.

The standard deviation of accuracy with the 16S rRNA was a significant 19.8 times higher (BLAST) and 20.7 times higher (k-mer matching) compared to the rRNA operon. Additionally, the minimum accuracy observed with the rRNA operon consistently exceeded 0.95, whereas the 16S rRNA dipped below 0.5 in some cases.

### Microbial community composition prediction

We conducted simulations to evaluate the effectiveness of different regions for predicting the species compositions (Fig. 2). These simulations assumed species existed in random proportions following a Dirichlet distribution. The figure depicts the predicted proportions of the top 10 species for each method (rRNA operon, 16S rRNA, and 16S rRNA V3-V4 regions). Predictions using the rRNA operon closely matched the actual compositions. The 16S rRNA predictions displayed a similar trend to the actual compositions, but with some discrepancies in the ratios between species. Predictions based on the 16S rRNA V3-V4 regions deviated significantly from the actual compositions.

**FIG 2.**
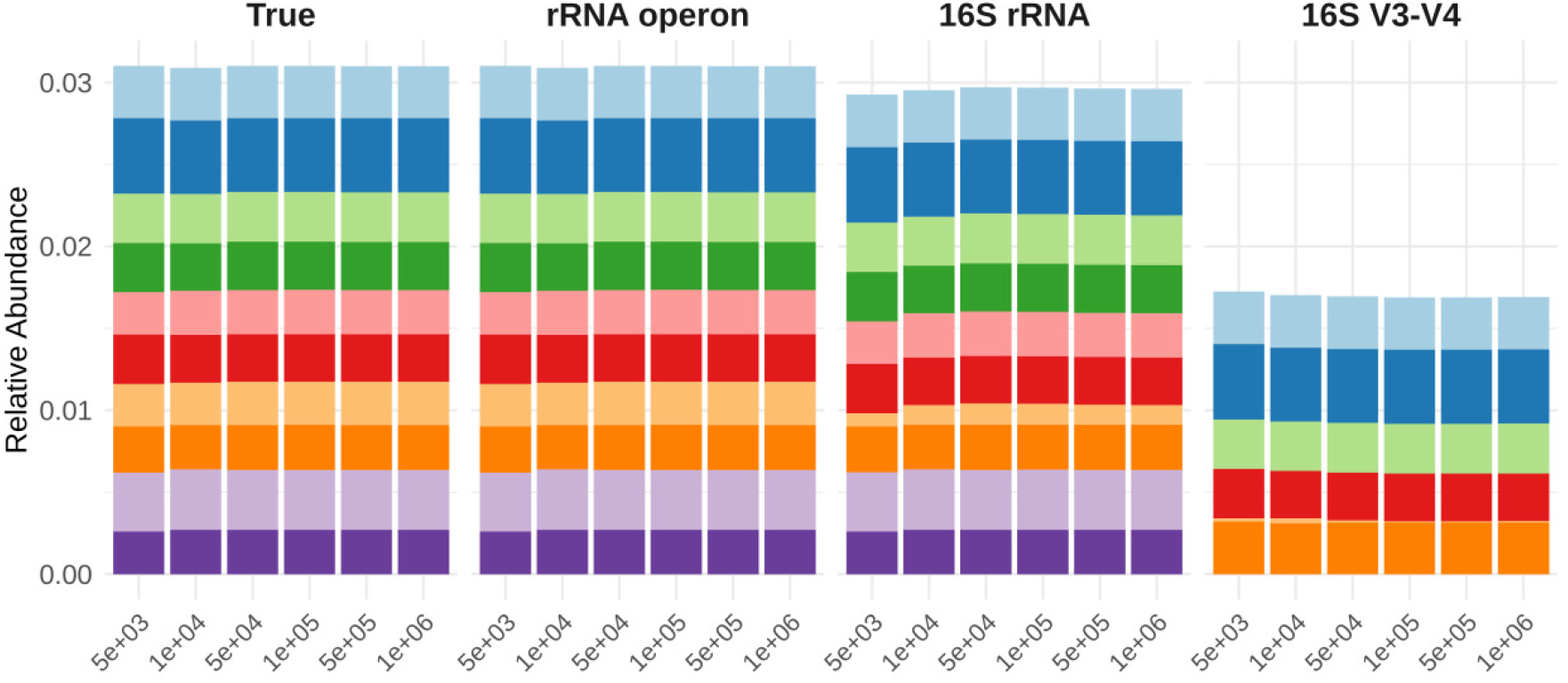
The relative abundance of the top 10 abundant species in the “True” data, where “True” represents the actual proportions. “rRNA operon,” “16S rRNA,” and “16S V3-V4” show the proportions of species predicted based on the accuracy of species classification within those genomic regions. Each color represents the same species across different predictions, with the x-axis indicating the number of reads used in the simulation and the y-axis showing the proportion of each species.

To numerically verify these observed trends, we calculated the Pearson correlation coefficient between the actual and predicted proportions (Table 1). Across six simulations, the correlation between actual and predicted proportions using the rRNA operon remained consistently high, with an average of 0.999 and a standard deviation of 0.0001. This held true regardless of the number of reads used in the simulation. The 16S rRNA exhibited a lower average correlation (0.849) with a higher standard deviation (0.014), indicating a poorer match to the actual proportions and greater variability between simulations compared to the rRNA operon. The 16S rRNA V3-V4 regions yielded the lowest average correlation (0.288) with a standard deviation of 0.022. The correlation between actual and predicted proportions increased with the number of simulated reads for both the 16S rRNA and its V3-V4 regions, reaching a plateau beyond 500,000 reads.

**TABLE 1.** Pearson correlations between actual species proportions and predicted species proportions using the rRNA operon, 16S rRNA, and 16S rRNA V3-V4 region, by the number of reads used for the simulation.

|  | **5,000** | **10,000** | **50,000** | **100,000** | **500,000** | **1,000,000** |
| --- | --- | --- | --- | --- | --- | --- |
| **rRNA operon** | 0.998 | 0.998 | 0.998 | 0.998 | 0.998 | 0.998 |
| **16S rRNA** | 0.804 | 0.821 | 0.835 | 0.837 | 0.838 | 0.838 |
| **16S V3-V4** | 0.295 | 0.321 | 0.344 | 0.346 | 0.348 | 0.348 |

To quantify the similarity between the actual and predicted microbial community compositions, we employed the Bray-Curtis distance metric. A smaller Bray-Curtis distance signifies greater similarity between the two datasets. The predicted composition using the rRNA operon yielded a remarkably close distance to the actual composition, averaging only 0.001 across six simulations. Conversely, the average distances observed when utilizing the 16S rRNA and 16S rRNA V3-V4 regions were higher, at 0.088 and 0.367 respectively. Predictions based on the rRNA operon exhibited the closest match to the actual community composition, with distances 71.2 times smaller than those obtained using the 16S rRNA.

To statistically validate these observations, we conducted an Analysis of Similarities (ANOSIM) test using Bray-Curtis distance. This non-parametric method evaluates the probability of observed differences in similarity between groups arising by chance. The results confirmed these findings. Predictions made with the rRNA operon yielded a p-value of 0.272, indicating no statistically significant difference from the actual community composition. Conversely, predictions utilizing the 16S rRNA and 16S rRNA V3-V4 regions produced p-values of 0.006 and 0.004, respectively. These significant p-values (p < 0.01) demonstrate that these methods yielded compositions statistically distinct from the actual community.

### Microbial community composition and differential abundance in human gut microbiome data

We evaluated the performance using the rRNA operon, 16S rRNA, and 16S rRNA V3-V4 regions for microbal community composition prediction using real human gut microbiome data (Fig. 3). The analysis included 558 overlapping species from samples of 14 healthy donors and 14 patients. The correlation between the reference species proportions and those predicted using the rRNA operon was remarkably high, averaging 1.00 with a minimal standard deviation of 0.000003. Predictions based on the 16S rRNA and the 16S V3-V4 regions exhibited lower correlations with the reference, with averages of 0.931 and 0.660, and standard deviations of 0.104 and 0.323, respectively. The rRNA operon consistently achieved high correlations between predicted proportions and reference proportions, with the lowest value still exceeding 0.999. Conversely, the 16S rRNA exhibited a significantly lower correlation, dropping as low as 0.620. These results align with our observations from the randomly generated data.

**FIG 3.**
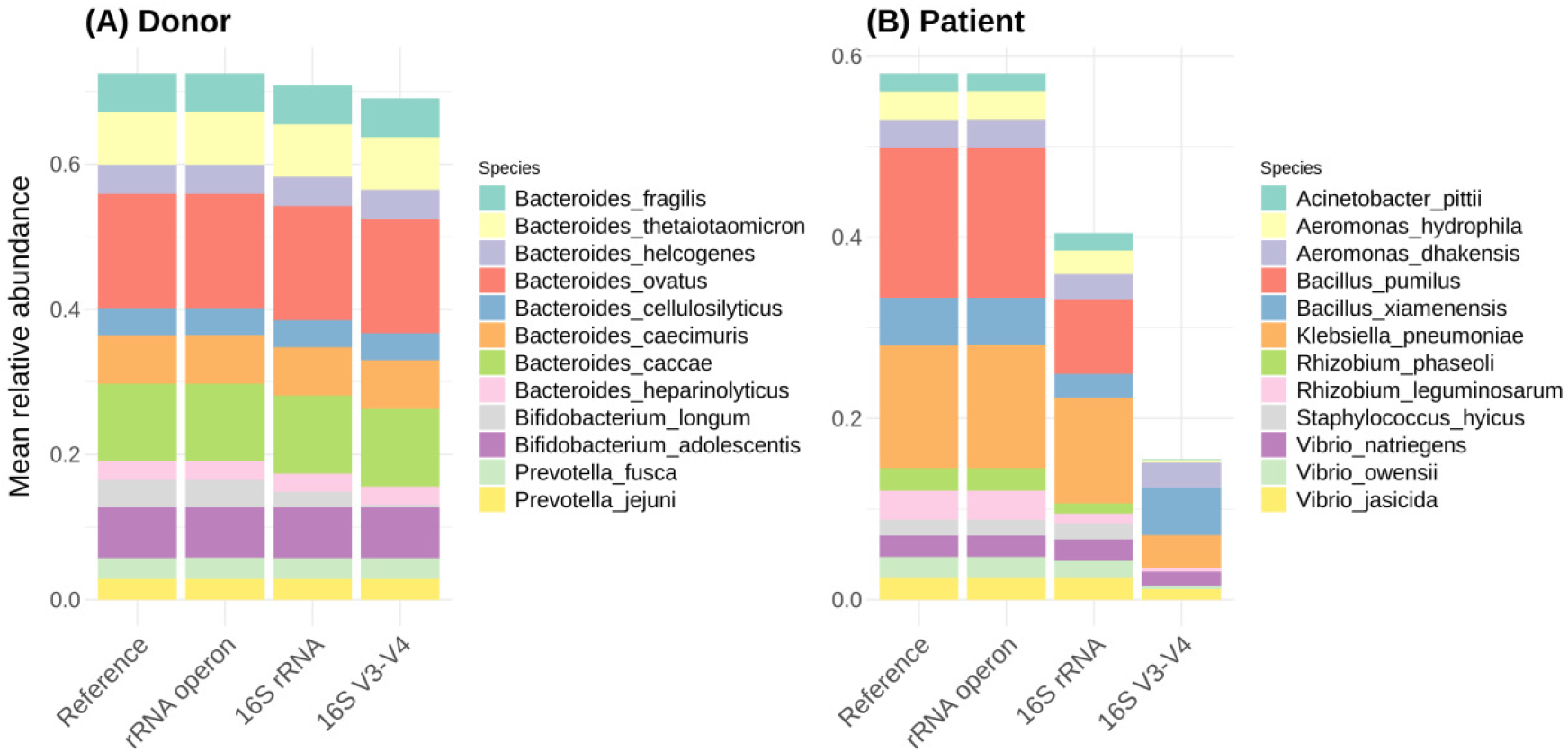
The relative abundance of the top 10 gut microbiome species in (A) FMT donors and (B) patients (B). “Reference” indicates actual proportions. “rRNA operon,” “16S rRNA,” and “16S V3-V4” show predicted proportions based on species classification accuracy.

We further assessed the methods by conducting differential abundance analyses based on both the reference compositions and those predicted by each classification method. We compared species identified as significantly different in each case (Fig. 4). The reference data identified 132 significantly differentially abundant species, which were accurately reflected by the predictions made with the rRNA operon. The 16S rRNA identified 151 significant species, with 127 overlapping with the reference findings. It missed 5 significant species (false negatives) and identified 24 species as significant that were not truly so (false positives). This translates to a false negative rate of 3.79% and a false positive rate of 18.2%, highlighting a higher prevalence of false positives with 16S rRNA. The 16S rRNA V3-V4 regions performed even worse, with even greater false negative (22.0%) and false positive (25.8%) rates.

**FIG 4.**
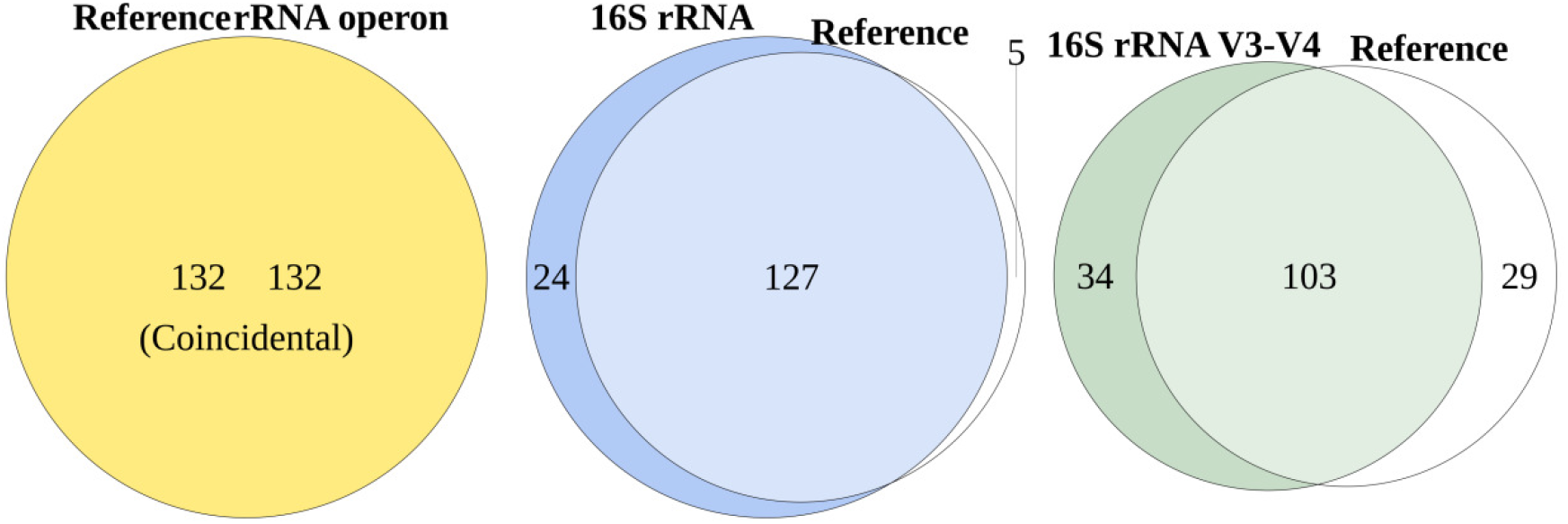
Venn diagrams of the significant species identified through differential abundance analysis based on proportions derived from the reference and rRNA operon, reference and 16S rRNA, and reference and16S rRNA V3-V4 regions. Overlapping sections of the diagram represent the number of species significantly identified across both methods. The area exclusive to the “Reference” (left side) shows the number of species that were false negatives. Areas unique to each method (right side) indicate false positives. Complete overlap between the “Reference” and a method implies identical species significance findings.

Fig. 5 depicts the coefficients of species that were identified as false negatives or false positives when using the 16S rRNA, as well as those whose coefficients differed in direction compared to using the reference. Here, the coefficient represents the relative abundance of a species in patients compared to donors. Some species identified as significantly different by 16S rRNA that were not detected by the reference or the rRNA operon predictions. Additionally, one species exhibited an opposing abundance trend between donors and patients when comparing the reference and 16S rRNA data. Furthermore, the magnitudes of the coefficients often differed between the reference and 16S rRNA findings. Conversely, the results using the rRNA operon displayed a high degree of agreement with the reference data, with consistent signs and magnitudes of coefficients for most species. Only two species showed minor discrepancies.

**FIG 5.**
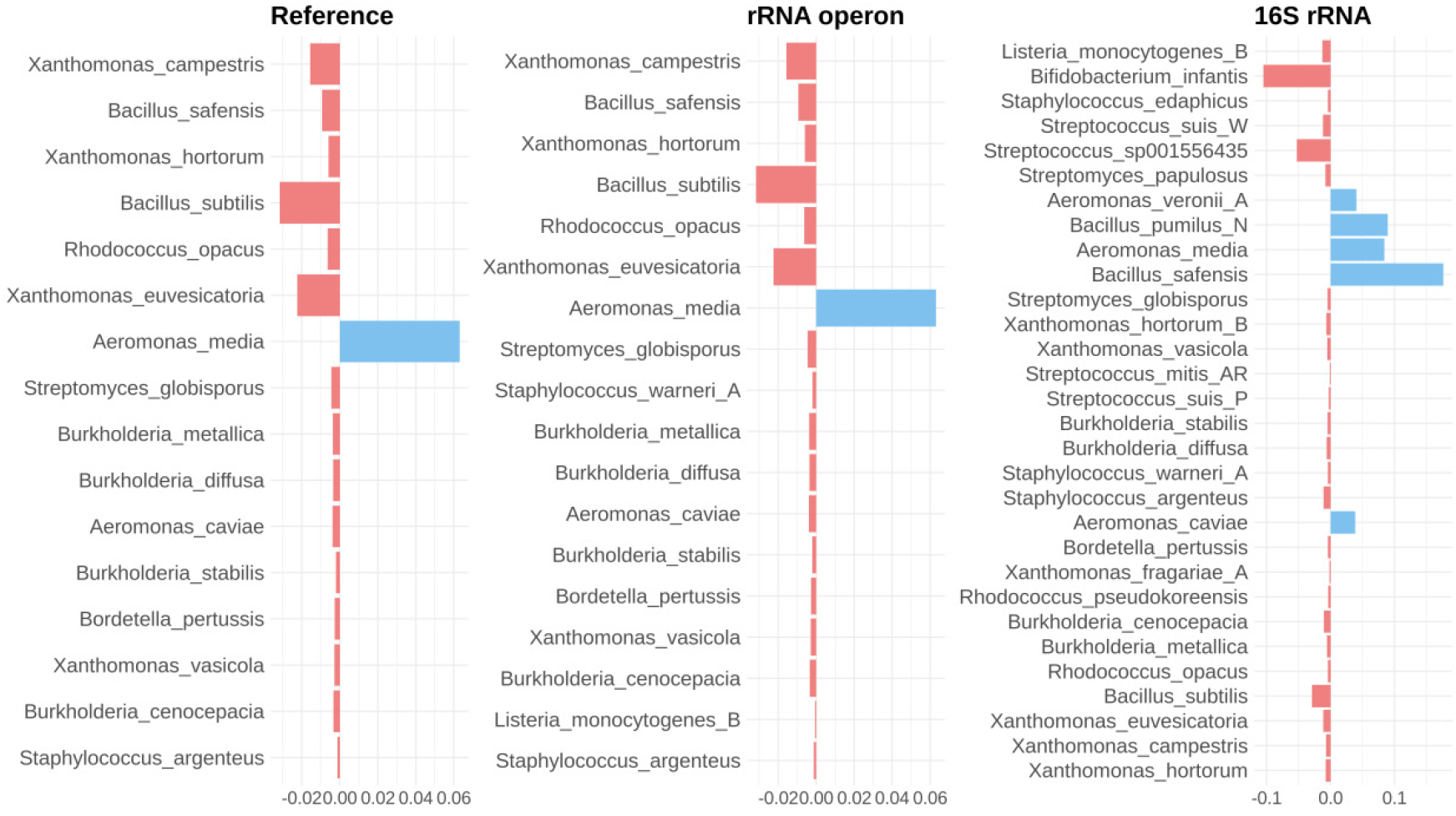
The coefficients from differential abundance analyses using the proportions obtained from reference, rRNA operon, and 16S rRNA. We only showed species that have discrepancy in the reference and 16S rRNA results; species identified as significant in one analysis but not the other and Species with differing direction of coefficient. A positive coefficient (depicted in pink) indicates a species is more abundant in patients, while a negative coefficient (shown in sky blue) suggests it is less abundant. The magnitude of the coefficient signifies the degree of abundance difference.

## DISCUSSION

The rRNA operon demonstrably outperformed the 16S rRNA gene in terms of species classification accuracy. Statistical tests confirmed this observation. Paired Wilcoxon rank sum tests revealed highly significant differences (p < 0.0001 for BLAST and k-mer matching) in accuracy favoring the rRNA operon. Furthermore, the rRNA operon exhibited considerably lower variability in accuracy across genera. This signifies that the rRNA operon offers consistently high and stable classification accuracy across various genera, while the 16S rRNA can yield unreliable results due to substantial variations in accuracy depending on the genus.

Simulations revealed a clear advantage for the rRNA operon in predicting species compositions within microbial communities. The correlation between actual and predicted proportions using the rRNA operon consistently outperformed both the 16S rRNA and the 16S rRNA V3-V4 regions. Notably, the correlation with the V3-V4 regions was significantly lower, rendering it unreliable for capturing meaningful relationships with the actual data. When using the rRNA operon versus the 16S rRNA, the difference in correlation between actual and predicted compositions in the data was greater than the difference in accuracy of individual species predictions. Similarly, the Bray-Curtis distance metric further supported the superiority of the rRNA operon. This suggests a far more accurate reflection of the true community structure when employing the rRNA operon compared to the 16S rRNA.

Simulations replicating the composition of actual human gut microbiomes yielded consistent results. Notably, this lower accuracy in predicting microbial compositions using 16S rRNA was particularly problematic for patient groups. In the patient data, the average correlation between predicted and reference compositions for the rRNA operon remained at 1.00, while the 16S rRNA only achieved an average of 0.870.

The observed difference in accuracy for community composition predictions also impacted the results of differential abundance analyses. When utilizing proportions derived from the rRNA operon, the analysis identified the same 132 significant species as those identified using the reference proportions, indicating perfect agreement. In contrast, the analysis based on 16S rRNA data and 16S rRNA V3-V4 region data yielded discrepancies, further solidifying the limitations of these methods for accurate prediction.

Using the rRNA operon as a marker provided higher accuracy in individual species classification than using the 16S rRNA or its V3-V4 regions, leading to more accurate community composition predictions and more reliable results in differential abundance analyses. However, sequencing costs may increase with the breadth of the region being read (24). Consequently, the choice of method should consider the required resolution and available budget. The 16S rRNA can be a suitable option when less precision is acceptable or species-level analysis is not necessary and genus-level identification suffices. On the other hand, for research requiring precise species-level analysis, such as discovering biomarkers, utilizing the microbiome for treatments, or other studies necessitating accurate species identification, the rRNA operon is preferable. This is especially true for disease-related microbial community studies, as the accuracy difference in community composition predictions between methods was more pronounced in patient groups, highlighting the importance of using the rRNA operon for more precise species differentiation in such contexts.

The accuracy of microbal community composition prediction using the 16S rRNA or its V3-V4 regions improves when using more number of reads, which mean sequencing more data. However, this also raises data production costs and should be carefully weighed. Given the same budget, producing less data with the rRNA operon may be more efficient than generating more data with the 16S rRNA.

Employing the rRNA operon as a marker demonstrably enhances individual species classification accuracy compared to the 16S rRNA gene. This translates to more precise predictions of microbial community compositions and more reliable differential abundance analysis results. The 16S rRNA V3-V4 region exhibited even lower accuracy across all scenarios compared to the full 16S rRNA, highlighting a significant decline in precision. Therefore, for research requiring accurate species classification, employing the rRNA operon as a marker appears to be the most appropriate choice. In microbial community studies aiming for precise species-level analysis, utilizing the rRNA operon is advisable as using the 16S rRNA has its limitations, and relying solely on its V3-V4 regions may make it challenging to achieve meaningful results.

## MATERIALS AND METHODS

### Data collection

We collected complete bacterial genomes available in the NCBI database as of November 29, 2023 (14). To ensure robust comparisons, we only considered genera containing more than 50 complete genomes. Our final dataset comprised 72 genera, 2,026 species, and 20,314 genome sequences (Supplementary Table 1).

### rRNA operon and 16S rRNA sequence extraction

We utilized riboSeed with its default settings to extract rRNA operon sequences (15). Following identification of rRNA gene regions using the riboscan command, we employed the riboselect command to locate rRNA operon regions containing 16S, 23S, and 5S rRNA. The corresponding rRNA operon sequences were then extracted. Implementing quality control, only sequences within the 4,000 to 6,000 base pair range were retained. 16S rRNA sequences were extracted from regions identified as 16S rRNA by the riboscan results.

We employed EMBOSS’s primer search tool to identify the V3-V4 regions within the previously extracted 16S rRNA sequences (16). The primer sequences used were ‘CCTACGGGNGGCWGCAG’ for the forward primer and ‘GACTACHVGGGTATCTAATCC’ for the reverse primer (17). A mismatch percentage of 10% was allowed during the search. Following quality control, only sequences with lengths between 430 bp and 550 bp were retained.

### Introduction of sequencing errors

To simulate real-world applications of using 16S rRNA and rRNA operon sequencing for species classification, we introduced random sequencing errors into the extracted sequences. Error rates were determined by referencing a 2022 study comparing the accuracy of Illumina and ONT technologies (18). 1D ONT MinION read error rates were applied to the rRNA operon sequences, while the average error rate of Illumina’s read1 and read2 was used for the 16S rRNA sequences (Table 2). Errors were introduced through random positional mismatches, insertions, and deletions.

**TABLE 2.** Error rates applied to simulate sequencing inaccuracies in rRNA operon and 16S rRNA sequences for species classification simulations. The table outlines the mismatch, insertion, and deletion rates for the rRNA operon sequenced with Nanopore technology and the 16S rRNA (including its V3-V4 regions) sequenced with Illumina.

| Marker region | Sequencing tool | Mismatch rate | Insertion rate | Deletion rate |
| --- | --- | --- | --- | --- |
| rRNA operon | Nanopore | 0.0116 | 0.0081 | 0.0144 |
| 16S rRNA | Illumina | 0.0089 | 0.00045 | 0.00045 |
| 16S rRNA V3-V4 | Illumina | 0.0089 | 0.00045 | 0.00045 |

For each position, a random number between 0 and 1 was generated. If this number was lower than the error rate, an error was introduced. Mismatches involved replacing the original nucleotide with a random one. Implementation was carried out using BioPython SeqIO (19).

### Species classification within genus

Two distinct methods were employed to classify species within the same genus: BLAST alignment score (20) and k-mer matching.

The BLAST-based approach used the sequences of the rRNA operon or 16S rRNA (including random errors) as the query, while the original sequences of the extracted regions served as the reference. Nucleotide BLAST was run with default options. Each sequence was classified into the species with the highest bitscore. In cases of ties, one species was randomly chosen for classification.

The k-mer matching method benchmarked the approach commonly used in microbiome data classification by Kraken (21). This method involves finding the number of exact matches of 31-mers and classifying the sequence to the species with the most 31-mer matches. Similar to the BLAST approach, ties were resolved by randomly choosing one species for classification.

In both methods, when multiple copies of the rRNA operon or 16S rRNA were present, we classified based on the copy with the highest score or the greatest number of matches. We assessed the accuracy of species classification per genus, by calculating the proportion of samples within each genus that were correctly assigned to their respective species.

### Simulation in microbial community data

To evaluate the accuracy of species classification in community data, a simulation was run on community composition data. First, we used a Dirichlet distribution to randomly set proportions for the species in our study and made a mock proportion data. Reads were initially distributed to match these true species proportions. Based on the likelihood of their classification through k-mer matching from the previous analysis, reads were then assigned to species. For example, if species A was correctly classified 90% of the time and misclassified as species B 10% of the time, a read intended for species A would have a 90% chance of being assigned to A and a 10% chance to B.

This process was applied to all reads. We conducted simulations for library sizes of 5,000, 10,000, 50,000, 100,000, 500,000, 1,000,000 reads.

### Microbial community analysis and differential abundance analysis in real world data

To further validate our findings using real-world gut microbiome data, we performed simulations with publicly available metagenomic proportions. We leveraged gut microbiome data from both donors and recipients of fecal microbiota transplantation (FMT) described in (22). Assuming the reported proportions reflect reality, we predicted species proportions based on the classification accuracy of the rRNA operon, 16S rRNA, and 16S rRNA V3-V4 regions. This procedure mirrored our previous community composition simulation, again using a library size of 100,000 reads. Subsequently, we employed the R package ‘Maaslin2’ to conduct differential abundance analysis comparing successful donors and pre-FMT patients (23). The analysis utilized AST transformation, TSS normalization, and a linear model. We considered findings with an adjusted false discovery rate (FDR) of less than 0.01 to be statistically significant.

## ACKNOWLEDGMENT

We would like to express our gratitude to eGnome Inc. for their support throughout the course of this research.

